# Therapeutic potential of oxygen through M101 transportation in reducing vaso-occlusion and hemolysis in a mouse model of sickle cell disease

**DOI:** 10.1101/2025.06.18.660442

**Authors:** Franck Zal, Rachel Rignault-Bricard, Leïla Demini, Benoît Barrou, Laurent Lantieri, Olivier Hermine, Jean-Benoît Arlet, Thiago Trovati Maciel

## Abstract

Sickle cell disease (SCD) is a genetic disorder marked by hemoglobin S–induced red blood cell (RBC) sickling, causing hemolysis, vaso-occlusion, and organ damage. Despite its severity, treatment options remain limited. RBC transfusion is a highly effective therapeutic option in SCD, particularly due to its capacity to improve oxygen delivery and reduce sickling by introducing healthy red blood cells, but it is associated with a significant risk of transfusion-related complications. Our study investigates the treatment potential of M101, a novel hemoglobin-based oxygen carrier derived from the marine worm *Arenicola marina*, in a mouse model of SCD. M101’s unique properties, including its high oxygen-carrying capacity and indirect antioxidant outcome, make it a promising candidate for ameliorating SCD pathophysiology. The study demonstrates that M101 administration significantly reduces RBC sickling and hemolysis in SCD mice. Furthermore, M101 infusion significantly prevents microvascular occlusion in liver and lung and attenuates the inflammatory response after vaso-occlusion induction. These findings highlight M101 strong potential as a novel therapeutic strategy for managing or preventing painful vaso-occlusive crises in SCD. Further research is warranted to determine the optimal dosing regimen and to evaluate the long-term safety and efficacy of M101 in humans.

## Introduction

Sickle cell disease (SCD) is a severe and debilitating inherited blood disorder affecting millions globally, with a particularly high prevalence in sub-Saharan Africa, India, the Middle East, and among populations of African descent worldwide. SCD arises from a single nucleotide mutation in the hemoglobin beta (HBB) gene, which results in the production of hemoglobin S (HbS) ^1,2^. Under deoxygenated conditions, HbS undergoes polymerization, triggering red blood cell (RBC) deformation into a sickle-like shape. These sickled RBCs exhibit reduced deformability, leading to hemolysis, impaired microvascular blood flow, and vaso-occlusion ^3^. The resultant pathophysiological cascade manifests clinically as chronic hemolytic anemia, acute vaso-occlusive crises (VOC), end-organ damage, and heightened susceptibility to infections, all of which significantly impair quality of life and contribute to early mortality ^3-5^.

The pathophysiology of SCD is multifactorial, involving chronic hemolysis, inflammation, endothelial dysfunction, and oxidative stress. Hemolysis releases free unstable hemoglobin and heme into the bloodstream, triggering oxidative stress and nitric oxide depletion, which impair vascular homeostasis and promote vaso-occlusion ^6,7^. Furthermore, sickled RBCs and activated leukocytes contribute to the generation of inflammatory cytokines and adhesion molecules, which amplify endothelial adhesion and microvascular obstruction. This creates a vicious cycle of hypoxia, ischemia-reperfusion injury, and progressive tissue damage ^3^.

Despite advances in understanding SCD pathophysiology and implementing evidence-based therapies, including hydroxyurea, chronic transfusions, and recently approved disease-modifying agents, significant therapeutic gaps remain. Hydroxyurea, the first FDA-approved disease-modifying therapy for SCD, increases fetal hemoglobin (HbF) production and reduces RBC sickling. Its clinical efficacy is well established, making it an essential component of disease management. Nevertheless, it remains insufficient in preventing VOCs or organ damage in some patients ^5,8^.

Red blood cell transfusion remains the other cornerstone treatment for severe acute complications, such as stroke or acute chest syndrome, with a remarkable efficacy. It is also used as long-term therapy to prevent vaso-occlusive events ^9^. The exact mechanism of action of RBC transfusions in SCD is not fully understood even if it is use on patient, but it is thought to involve improved oxygen delivery and reduced sickling through the introduction of healthy red blood cells. Unfortunately, transfusions are frequently complicated by severe alloimmunization reactions, such as delayed hemolytic transfusion reactions (DHTR), which can make further transfusions challenging ^10^. Moreover, repeated transfusions may lead to secondary iron overload (hemochromatosis) and increase the risk of transfusion-transmitted infections, including viral pathogens and malaria, particularly in African settings ^11^.

Building on the proven efficacy of transfusion, hemoglobin-based oxygen carriers (HBOCs) are being explored as substitutes to reduce immunological risks and iron overload ^12,13^. However, earlier generations of HBOCs faced significant limitations, including oxidative toxicity, instability, and vasoactivity, which have restricted their clinical utility. Preclinical studies have highlighted the importance of developing HBOCs with intrinsic antioxidant properties to counteract the oxidative stress and inflammation that exacerbate SCD pathology^14^.

M101, a novel natural extracellular hemoglobin derived from the marine lugworm *Arenicola marina*, offers unique advantages over conventional HBOCs. Structurally and functionally distinct, M101 has a high oxygen-carrying capacity (managed by physical means according to partial pressure properties), stability across a broad range of environmental conditions, and intrinsic antioxidant properties conferred by copper/zinc superoxide dismutase-like (Cu/Zn-SOD) activity. These antioxidant properties and indirect anti-inflammatory attributes are particularly relevant in SCD, where oxidative stress contributes to hemolysis, endothelial injury, and VOC ^15,16^. Unlike previous HBOCs, M101 has demonstrated excellent biocompatibility, absence of vasoactivity, and the ability to maintain tissue oxygenation without causing vasoconstriction ^17,18^. Notably, M101’s antioxidant properties allow it to scavenge free radicals and reduce oxidative damage, which is a critical driver of RBC dysfunction and endothelial activation in SCD ^15^.

Preclinical studies in other hypoxia-driven models, such as ischemia-reperfusion, have shown that M101 can improve tissue oxygenation by physical mechanism, reducing oxidative stress and consequently protect against inflammatory injury, although M101 has no intrinsic anti-inflammatory properties ^19^. These findings suggest that M101’s multimodal mechanisms (oxygen delivery, oxidative stress mitigation and indirect anti-inflammatory effects) may have significant treatment potential in SCD, where these processes are central to disease pathophysiology. Preclinical studies in dogs and monkeys have demonstrated a good tolerability of the intravenous injections of M101 ^20^. In addition, M101 safety and efficacy was demonstrated in the field of human transplantation, preserving kidney, limbs and face before grafting ^21,22^.

In this study, we explored the therapeutic potential of M101 in a well-established mouse model of SCD. We specifically investigated the effects of M101 on RBC sickling, hemolysis, and microvascular occlusion in two relevant VOC models: hypoxia-induced and heme-induced vaso-occlusion. Our findings highlight the ability of M101 to mitigate key pathological features of SCD, underscoring its potential as a novel treatment targeting hypoxia and oxidative stress in SCD. By addressing these interconnected mechanisms, M101 represents a promising candidate for improving outcomes in patients with SCD, particularly during acute vaso-occlusive episodes where tissue hypoxia and oxidative stress are most severe.

## Methods

### Animal Care and Experimental Design

All animal experiments were conducted in compliance with the *European Directive 2010/63/EU*, the *Care and Use of Laboratory Animals Guidelines*. Ethical approval was obtained from the Institutional Animal Care and Use Committee (IACUC) and authorized by the French Ministry for Research (permit number 201809051638898). Mice were housed in ventilated cages (4–5 mice/cage) under controlled conditions (temperature: 22 ± 2°C; humidity: 50 ± 5%; 12-hour light-dark cycle) with ad libitum access to food and water.

Townes-HbSS mice (Jackson Laboratory, Bar Harbor, ME, USA) were used in this study ^23^. These mice, with a mixed 129/B6 genetic background, harbor deletions of the endogenous α- and β-globin loci replaced by human α- and AγβS globins. Homozygous βS/βS mice develop hallmark SCD features, including chronic anemia, hemolysis, and sickle-shaped RBCs. Mice were randomized into treatment groups to ensure balanced baseline characteristics across groups.

### Red Blood Cell Half-Life Analysis by Flow Cytometry

To measure RBC lifespan, HbSS mice were intravenously injected with 1 mg of biotin (EZ-Link Sulfo-NHS-Biotin, Life Technologies) and treated with either M101 or phosphate-buffered saline (PBS) as control. Blood samples (5 µL) were collected retro-orbitally on days 0, 1, 3, 6, and 9 post-treatment using EDTA-coated tubes to prevent clotting.

Biotin-labeled RBCs were stained with the following antibodies: anti-TER-119 conjugated to Pacific Blue (Biolegend, Clone TER-119) and streptavidin–Alexa Fluor 647 (Biolegend, 405237). Cells were also stained with Syto16 (Life Technologies) to exclude nucleated cells and reticulocytes. After 30 minutes of staining at 4°C, samples were washed with fluorescence-activated cell sorting (FACS) staining buffer and analyzed on a Gallios™ flow cytometer (Beckman Coulter). Post-acquisition analysis was performed using *FlowJo* software (v10.0; FlowJo LLC). The proportion of biotin-positive RBCs was used to calculate RBC survival over time.

### Induction of Vaso-Occlusion and Hemolysis

Townes HbSS mice were randomly assigned to receive either M101 or phosphate-buffered saline (PBS) as a control. Treatments were administered intravenously one hour prior to the induction of vaso-occlusion.

To model hypoxia-induced VOC, mice were subjected to hypoxia-reoxygenation stress using a BioSpherix hypoxia chamber equipped with a ProOx 110 gas controller. Animals were exposed to 8% O_2_ for 3 hours, followed by reoxygenation in ambient air (21% O_2_) for 1 hour.

For the heme-induced VOC model, free heme was introduced by intravenous injection of hemin (Sigma-Aldrich), freshly dissolved in 0.25 M NaOH, neutralized to pH 7.4 with PBS, and sterile-filtered through a 0.22 µm membrane. Hemin was administered at a dose of 50 µmol/kg. Mice were euthanized for analysis three hours post-injection.

Blood samples were collected under ketamine-xylazine anesthesia (80 and 10 mg/kg, respectively) via heparinized capillaries. Complete blood counts (CBCs) were performed using a ProCyte Dx Hematology Analyzer (IDEXX, France) in accordance with the manufacturer’s instructions.

### Hemolysis Biomarker Analysis

Plasma was isolated by centrifuging whole blood samples (15 min at 2000×g, 4°C) collected in K2 EDTA tubes (Melet Schloesing Laboratoires, France). Aliquots were stored at -80°C until analysis. The following biomarkers of hemolysis were analyzed from whole blood plasma using commercial assays: (1) total plasma hemin (TPH) measured using a Hemin Assay Kit (Sigma-Aldrich), (2) total bilirubin measured using a Bilirubin Assay Kit (Sigma-Aldrich), (3) lactate dehydrogenase (LDH) measured using a Pierce LDH Cytotoxicity Assay Kit (Thermo Fisher Scientific, Waltham, MA, USA).

### Red Blood Cell Sickling Analysis

Following hypoxia-reoxygenation exposure, whole blood samples from M101- and PBS-treated mice were collected and fixed in 0.5% glutaraldehyde. Fixed blood was diluted 1:100 in PBS, and the percentage of sickled RBCs was quantified by light microscopy. Images were acquired at 100× magnification, and five random fields per sample were analyzed.

### Quantification of Vaso-Occlusion by Immunofluorescence

At study termination, mice were euthanized by cervical dislocation. To assess vascular occlusion in target organs, mice were perfused with 1 mL saline solution through the left ventricle, and lungs and liver were collected, weighed, and processed for histology.

Tissue sections (5 µm) were deparaffinized, rehydrated, and subjected to antigen retrieval using citrate buffer (pH 6.0; Biolegend). Vaso-occluded areas were visualized by immunostaining with anti-TER-119 antibodies conjugated to Alexa Fluor-488 or Alexa Fluor-647 (1:100 dilution, Biolegend). Slides were mounted using ProLong Diamond Antifade Mountant with DAPI (Thermo Fisher Scientific).

Images were captured on an EVOS M5000 Imaging System (Thermo Fisher Scientific) at 200× magnification. Quantitative analysis of vaso-occluded regions was performed in *ImageJ* software. Results were expressed as the percentage of vaso-occluded area per field of view (FOV).

### Quantitative Real-Time PCR

Total RNA was extracted from frozen liver and lung tissues using the TissueRuptor homogenizer and the RNeasy Mini Kit (Qiagen) according to the manufacturer’s instructions. RNA concentration and purity were assessed using a NanoDrop spectrophotometer (Thermo Fisher Scientific). Complementary DNA (cDNA) was synthesized from 1 µg of total RNA using the iScript cDNA Synthesis Kit (Bio-Rad). Quantitative real-time PCR (qPCR) was performed using the SsoAdvanced Universal SYBR Green Supermix (Bio-Rad) on a CFX96 Touch Real-Time PCR Detection System (Bio-Rad). Reactions were carried out in technical duplicates, and gene expression levels were normalized to housekeeping genes Actb and Rpl4. Relative mRNA expression was calculated using the ΔΔCt method, and data analysis was performed using CFX Manager Software (Bio-Rad) with the regression mode enabled for quantification.

### Statistical Analysis

Data are presented as mean ± standard error of the mean (SEM). Statistical analyses were conducted using *GraphPad Prism* (v6.00 for in vitro and v9.00 for in vivo, GraphPad Software, San Diego, CA, USA). Group comparisons were made using one-way analysis of variance (ANOVA) followed by Tukey’s multiple comparisons test. A *P*-value <0.05 was considered statistically significant.

For all analyses, experimental replicates and sample sizes are reported in the figure legends.

## Results

### M101 Reduces Sickling in vivo and in vitro

To evaluate the anti-sickling properties of M101, we administered single intravenous doses of M101 (100, 300, 600, or 1200 mg/kg) to Townes HbSS mice, followed by exposure to a hypoxia challenge (8% O_2_ for 3 hours). Sickling of RBCs was quantified to determine the extent of deformation. M101 significantly decreased the percentage of sickled RBCs in a dose-dependent manner, demonstrating potent *in vivo* anti-sickling efficacy. Notably, even at the lowest dose (100 mg/kg), M101 reduced sickling to 25.0±0.1% compared to 38.0±0.8% in vehicle-treated controls (p<0.01), indicating robust inhibition of RBC sickling at physiologically relevant doses. The anti-sickling effect was further enhanced with increasing doses, achieving reductions to 17.0±1.2% at 600 mg/kg and 16.0±1.6% at 1200 mg/kg (both p<0.001 vs. control; Figure 1A). Based on these results, we therefore decided to evaluate M101 optimal dosing at 300 mg/kg and 600 mg/kg.

**Figure 1.**
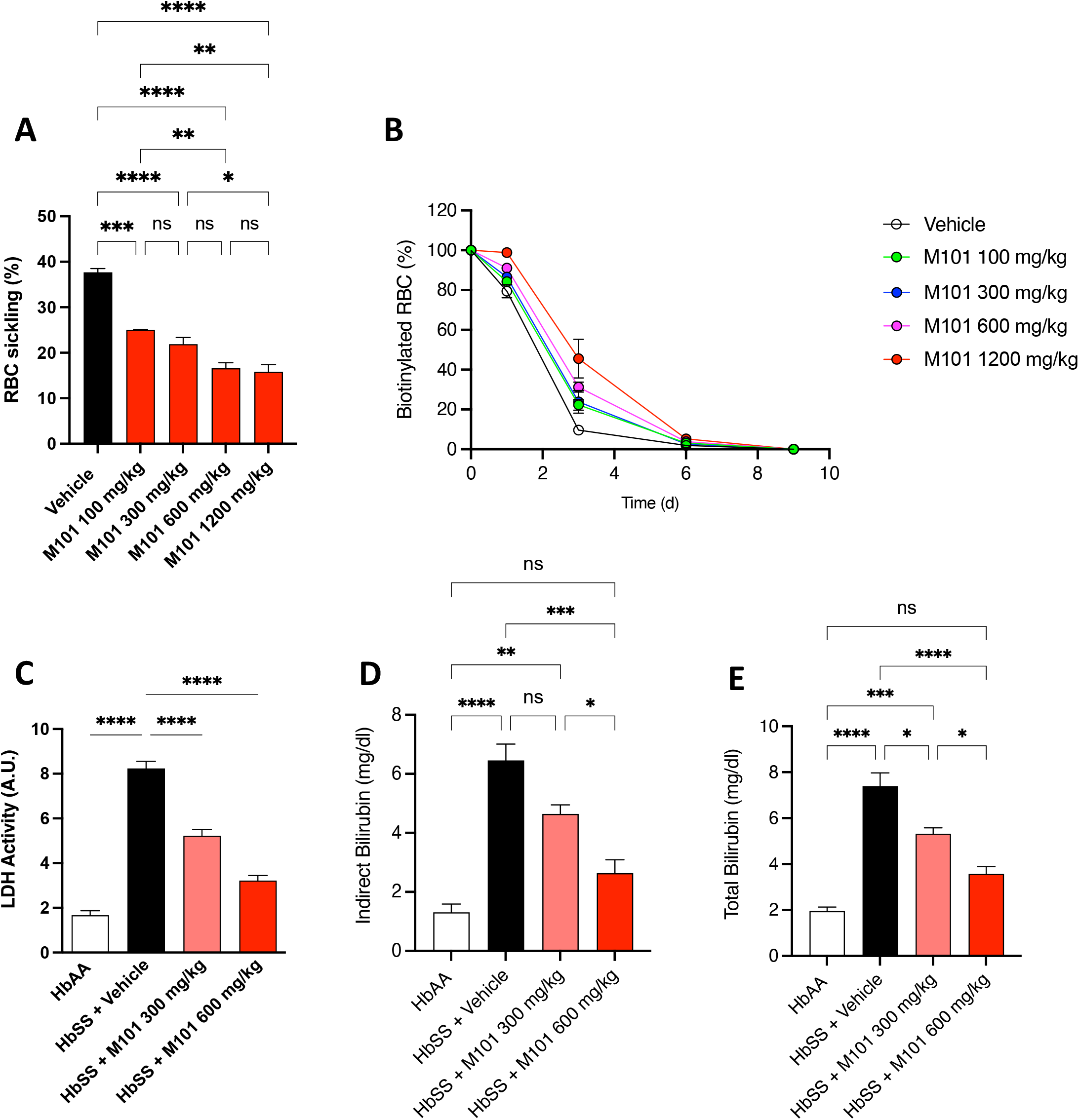
M101 reduces red blood cell sickling and prolongs RBC survival in hypoxia-challenged HbSS mice. (A) Percentage of sickled red blood cells (RBCs) in HbSS mice after hypoxia (8% O_2_, 3 h) with or without pretreatment with escalating doses of M101. (B) Biotin-labeled RBCs were tracked over time to assess RBC half-life in vivo following a single injection of M101 (100–1200 mg/kg). (C–E) Plasma biomarkers of hemolysis including total bilirubin (C), indirect bilirubin (D), and lactate dehydrogenase (LDH) activity (E) were quantified in HbSS mice treated with vehicle or M101 (300 or 600 mg/kg). HbAA mice served as healthy controls. Data are presented as mean ± SEM. Statistical significance determined by one-way ANOVA with Tukey’s post hoc test. p<0.05, p<0.01, p<0.001, p<0.0001.

### M101 Ameliorates Hemolysis and Prolongs RBC Survival

To investigate the effect of M101 on RBC survival, we performed biotinylation labeling in Townes HbSS mice, allowing tracking of RBC half-life following treatment. Vehicle-treated HbSS mice exhibited significantly shortened RBC survival. M101 treatment notably prolonged RBC half-life, indicating improved RBC stability (Figure 1B).

Markers of hemolysis, including serum LDH and bilirubin levels, were assessed in mice exposed to hypoxia-reoxygenation stress (3 hours hypoxia followed by 1 hour reoxygenation). M101 administration (300 and 600 mg/kg, 1 hour prior to hypoxia) resulted in dose-dependent reductions in serum LDH levels. Specifically, LDH was reduced by 37% at 300 mg/kg (5.22±0.28 AU vs. 8.24±0.31 AU in controls, p<0.01) and by 61% at 600 mg/kg (3.22±0.23 AU, p<0.01; Figure 1C).

Indirect bilirubin, another marker of hemolysis, was also significantly decreased by M101 treatment. M101 at 600 mg/kg reduced indirect bilirubin levels by 60% (2.6±0.45 mg/dl vs. 6.5±0.55 mg/dl in controls, p<0.05; Figure 1D). Total bilirubin was similarly modulated (Figure 1E), with reductions of 29% at 300 mg/kg (4.6±0.30 mg/dl, p<0.05) and 51% at 600 mg/kg (3.6±0.31 mg/dl, p<0.01), compared to vehicle-treated group (6.5±0.55 mg/dl). These findings provide evidences that M101 not only mitigates hemolysis but also improves RBC survival in HbSS mice subjected to hemolytic stress.

### M101 Improves Microvascular Occlusion Under Hypoxic and Heme-Induced Stress

Microvascular occlusion, a defining feature of VOC, arises from the obstruction of small blood vessels by sickled RBCs. To evaluate the impact of M101 on microvascular congestion, we performed immunofluorescence staining using Ter-119, an erythroid marker, to quantify RBC accumulation in the liver and lungs of Townes HbSS mice.

In the hypoxia-reoxygenation model, vehicle-treated HbSS mice exhibited pronounced vascular congestion (Figure 2), with obstructed areas accounting for 25.6 ± 3.23% of the FOV in the lungs and 20.4 ± 0.56% in the liver. Pretreatment with M101 at 300 mg/kg significantly reduced vascular occlusion in both organs, decreasing obstruction to 15.6 ± 2.03% in the lungs (p < 0.05) and 9.5 ± 0.61% in the liver (p < 0.01). A higher dose of 600 mg/kg further enhanced this effect, reducing occlusion to 7.5 ± 0.55% in the lungs and 4.2 ± 0.25% in the liver (both p < 0.01 vs. vehicle).

**Figure 2.**
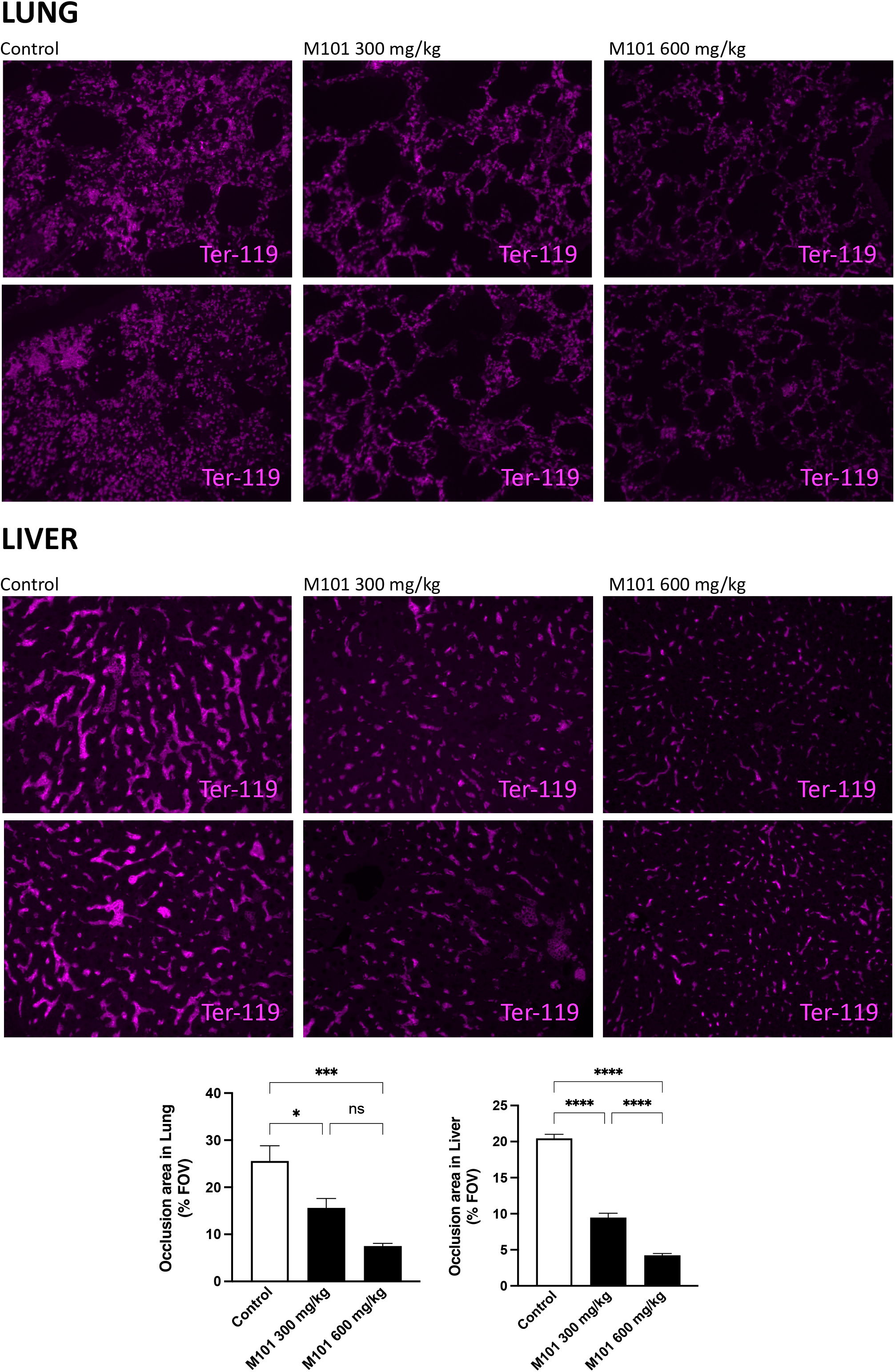
M101 ameliorates microvascular occlusion in liver and lung under hypoxic conditions. Quantification of Ter-119+ RBC-occluded areas in (A) liver and (B) lung tissues of HbSS mice following hypoxia-reoxygenation (8% O_2_ for 3 h followed by 1 h reoxygenation). Mice were pretreated with vehicle, M101 300 mg/kg, or M101 600 mg/kg 1 h prior to hypoxia. Occlusion is expressed as percentage of the field of view (FOV) occupied by congested vessels. Data are mean ± SEM. p<0.05, p<0.01, p<0.001, p<0.0001 by one-way ANOVA.

To evaluate M101 in a second model of VOC, mice were challenged with hemin, which induces microvascular congestion through oxidative stress and endothelial injury. In vehicle-treated animals, heme exposure resulted in occlusion of 23.5 ± 0.97% of the FOV in the lungs and 22.3 ± 0.78% in the liver. Treatment with M101 at 600 mg/kg significantly mitigated this heme-induced vascular congestion, reducing obstruction to 12.3 ± 0.41% in the lungs and 13.5 ± 0.93% in the liver (both p < 0.01 vs. vehicle; Figure 3), while the 300 mg/kg dose induces a moderate, non-significant decrease.

**Figure 3.**
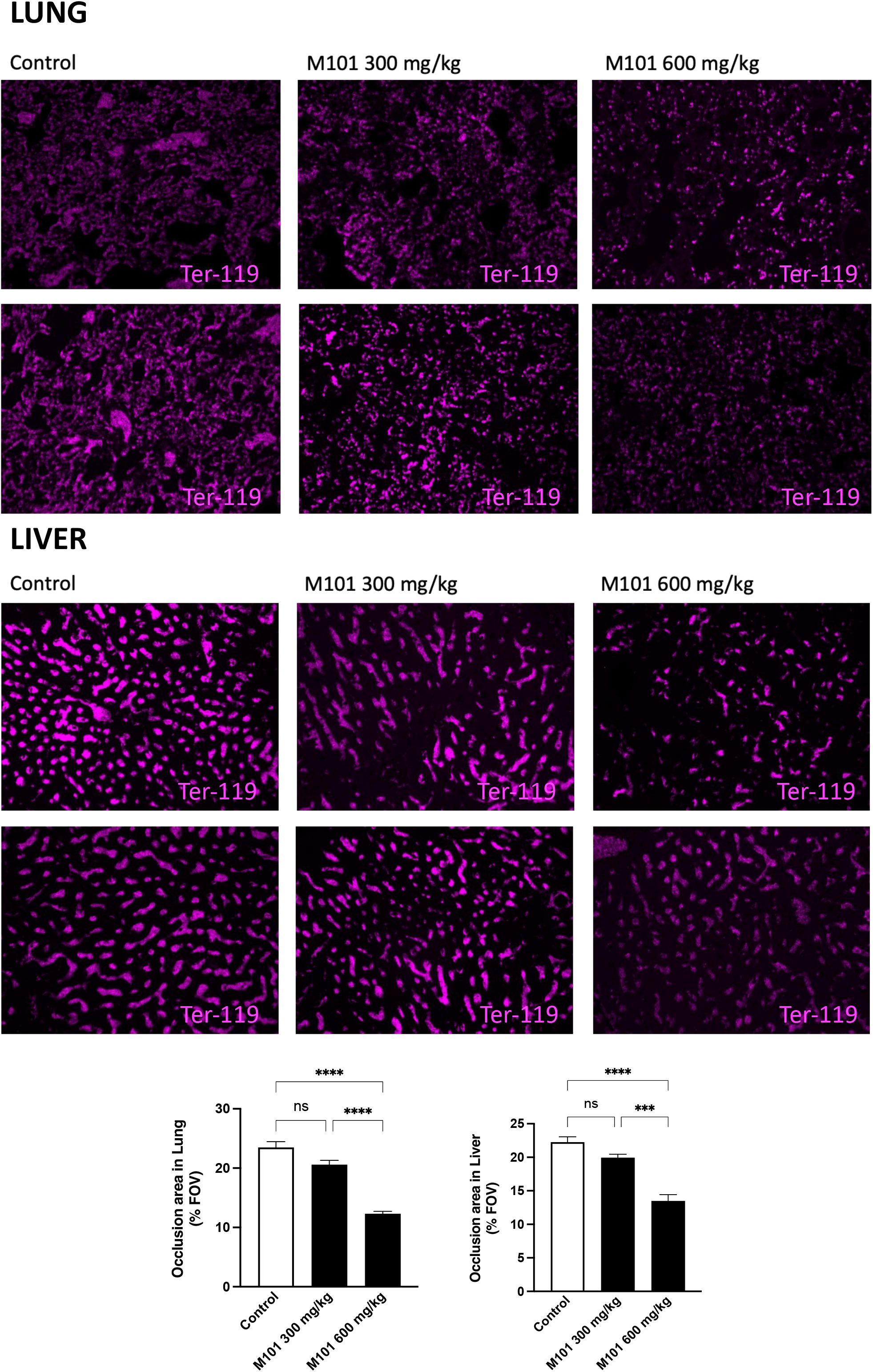
M101 reduces hemin-induced vascular congestion in liver and lung. Quantification of RBC occlusion in (A) liver and (B) lung of HbSS mice 3 h after intravenous hemin challenge (50 µmol/kg). Animals were pretreated with vehicle or M101 (300 or 600 mg/kg) 1 h before heme injection. Vascular occlusion was assessed by Ter-119 immunostaining and is expressed as a percentage of the field of view (FOV). Data are mean ± SEM. p<0.05, p<0.01, p<0.001, p<0.0001.

Together, these findings demonstrate that M101 effectively improves microvascular perfusion and alleviates vessel obstruction in both hypoxia and heme-induced models of VOC, underscoring its therapeutic potential in targeting microvascular dysfunction in SCD.

### M101 Suppresses Inflammatory Gene Expression in Liver and Lung Tissues Under Hypoxia and Hemin-Induced Vaso-Occlusive Stress

To evaluate the indirect anti-inflammatory effects of M101 during VOC, we performed quantitative real-time PCR to assess the expression of key inflammatory and oxidative stress-related genes in liver and lung tissues from HbSS mice subjected to either hypoxia-reoxygenation or hemin challenge.

In the hypoxia model, HbSS mice exhibited substantial upregulation of multiple pro-inflammatory cytokines and interferons in both organs. In the liver (Figure 4), Il1b expression increased approximately 4-fold compared to HbAA controls and was reduced by nearly 50% following M101 treatment at 600 mg/kg (p<0.05). Il6 levels were elevated nearly 150-fold and were suppressed by more than 60% with M101 (p<0.01), while Tnfa rose approximately 5-fold and was similarly decreased by over 50% (p<0.01). Interferon-related genes including Ifng, Ifna, and Ifnb1 were also significantly elevated, with M101 treatment resulting in reductions of more than 50% (p<0.05), indicating a broad indirect anti-inflammatory effect.

**Figure 4.**
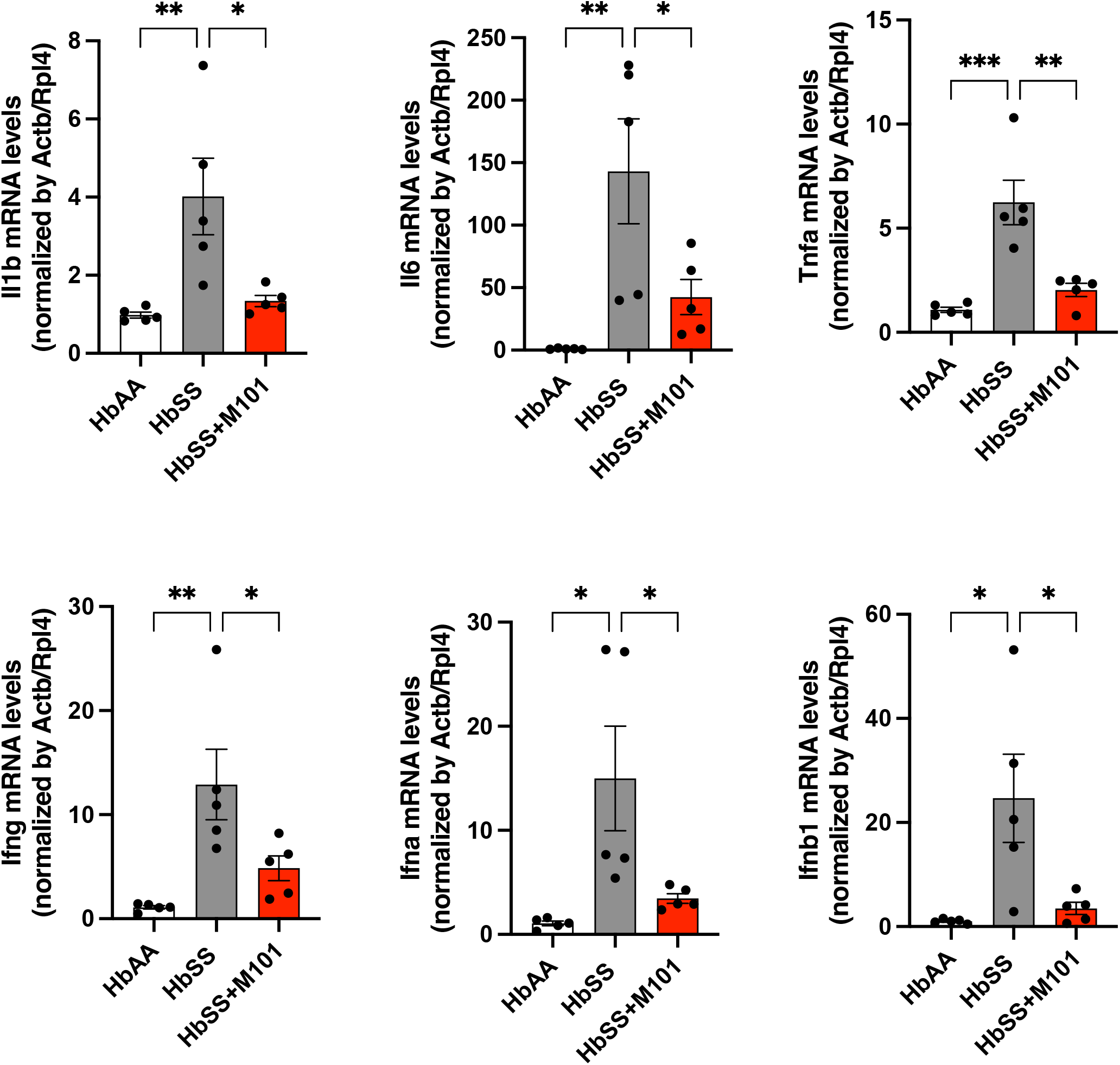
M101 suppresses indirectly hypoxia-induced inflammatory gene expression in the liver. Quantitative RT-PCR analysis of inflammatory cytokine and interferon transcripts in liver tissue from HbSS mice subjected to hypoxia-reoxygenation with or without M101 pretreatment (600 mg/kg). mRNA levels of Il1b, Ifng, Il6, Ifna, Tnfa, and Ifnb1 were normalized to Actb/Rpl4 and are shown relative to HbAA controls. Bars represent mean ± SEM. p<0.05, p<0.01, p<0.001 by one-way ANOVA.

In the lungs (Figure 5), hypoxia induced a 5-fold increase in Il1b, a ∼6-fold rise in Il6, and a 2-fold upregulation of Tnfa, all of which were significantly decreased with M101 (p<0.01). Ifnb1 expression was also reduced by more than 60% (p<0.01), while Ifng expression showed a more modest decrease (∼30%, p<0.05). Interestingly, Ifna levels in the lung were not significantly altered under hypoxia, with or without M101.

**Figure 5.**
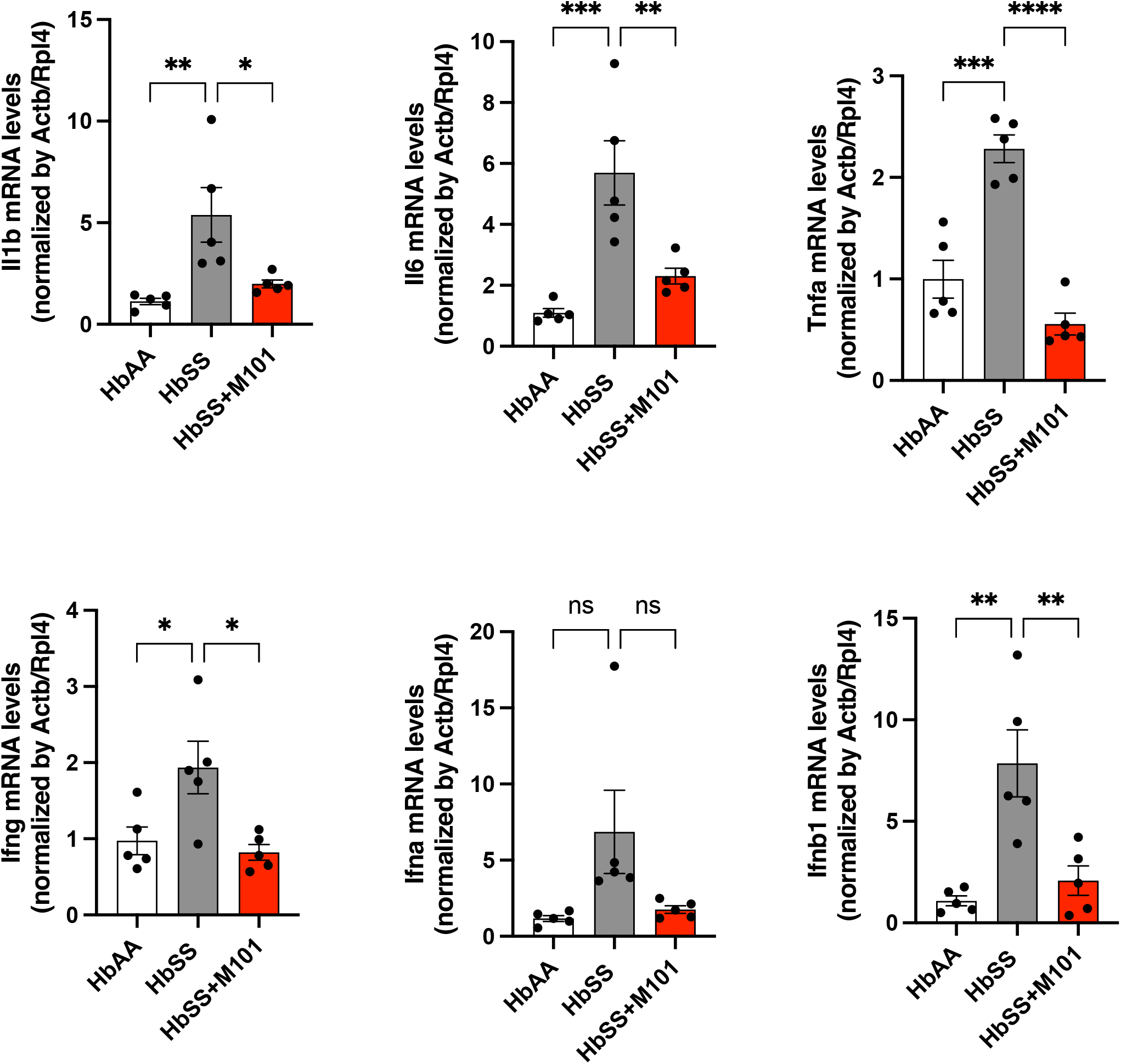
M101 avoids hypoxia-induced inflammatory responses in the lung. mRNA expression of pro-inflammatory cytokines and interferons in lung tissue of HbSS mice exposed to hypoxia-reoxygenation. Treatment with M101 (600 mg/kg) resulted in significant suppression of Il1b, Ifng, Il6, Tnfa, and Ifnb1. Ifna levels were not significantly altered. Gene expression was normalized to Actb/Rpl4. Data are presented as mean ± SEM. p<0.05, p<0.01, p<0.001, p<0.0001.

In the hemin-induced VOC model, a similarly robust inflammatory transcriptional response was observed. In the liver (Figure 6), hemin exposure resulted in significant increases in Il1b, Il6, and Tnfa, which were reduced following M101 administration (p<0.05 for all). The oxidative stress marker Hmox1 was elevated approximately 25-fold and was suppressed by over 50% with M101 (p<0.001). Moreover, interferon genes Ifng and Ifnb1 were significantly downregulated in M101-treated animals (p<0.05). In the lungs of hemin-challenged mice (Figure 7), expression of Il1b, Il6, and Tnfa was attenuated by M101 by more than 50% (p<0.05), while Hmox1 was suppressed by more than 60% from a ∼6-fold induction (p<0.01). Ifna and Ifnb1 levels were significantly reduced by M101 (p<0.05), whereas Ifng remained largely unchanged. Together, these findings demonstrate that M101 exerts potent indirect anti-inflammatory and anti-oxidative effects in both liver and lung tissues during VOC, irrespective of the underlying trigger. By significantly attenuating the expression of inflammatory cytokines and oxidative stress markers, M101 further supports its treatment potential as a systemic modulator of VOC-associated inflammation in sickle cell disease.

**Figure 6.**
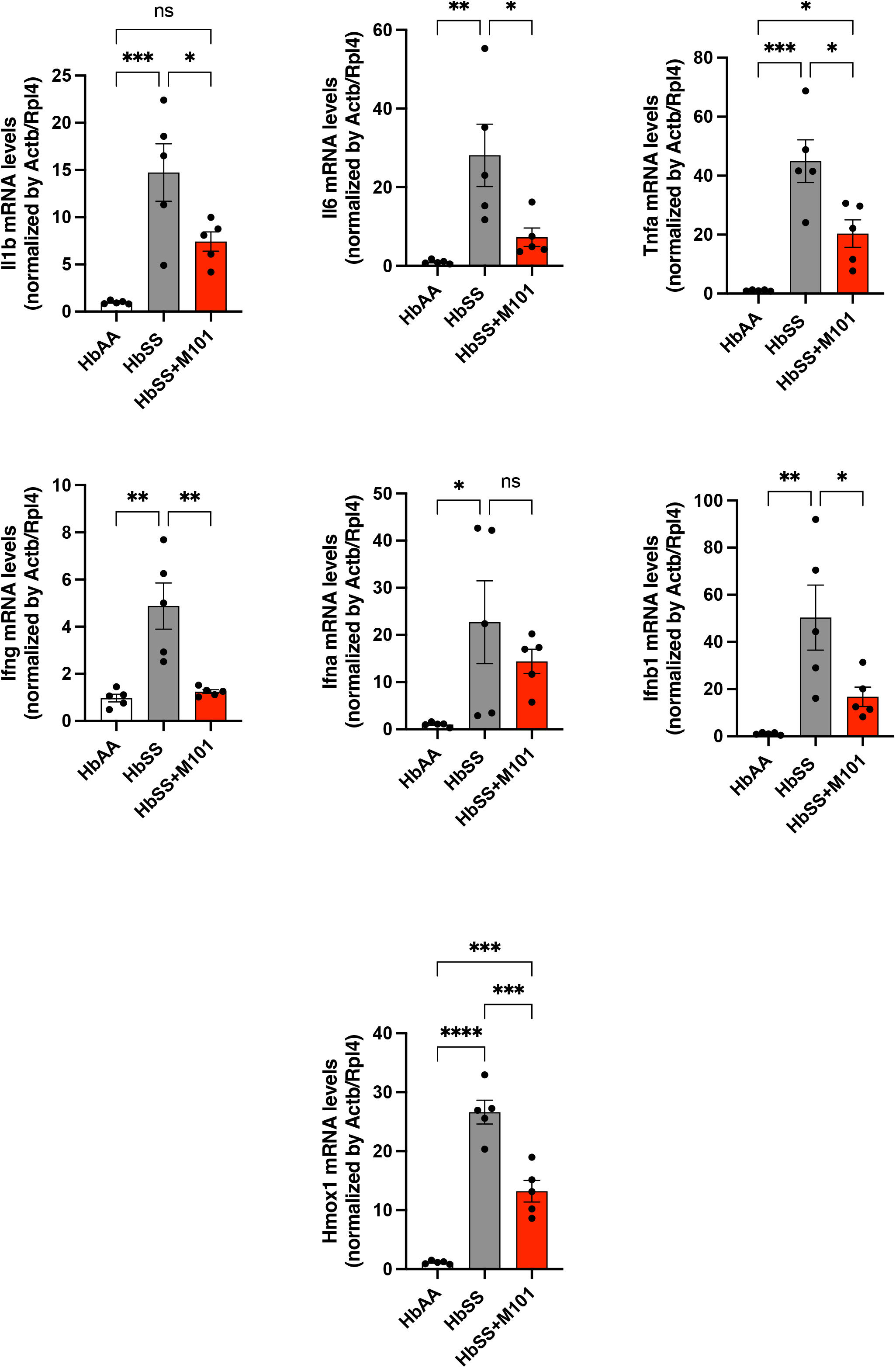
M101 attenuates inflammatory gene expression in the liver following hemin-induced stress. Liver mRNA expression of inflammatory (Il1b, Il6, Tnfa) and interferon-related genes (Ifng, Ifna, Ifnb1), as well as the oxidative stress marker Hmox1, following hemin administration (50 µmol/kg). M101 (600 mg/kg) significantly reduced gene expression in HbSS mice relative to vehicle-treated controls. Data normalized to Actb/Rpl4 and shown as mean ± SEM. p<0.05, p<0.01, p<0.001, p<0.0001.

**Figure 7.**
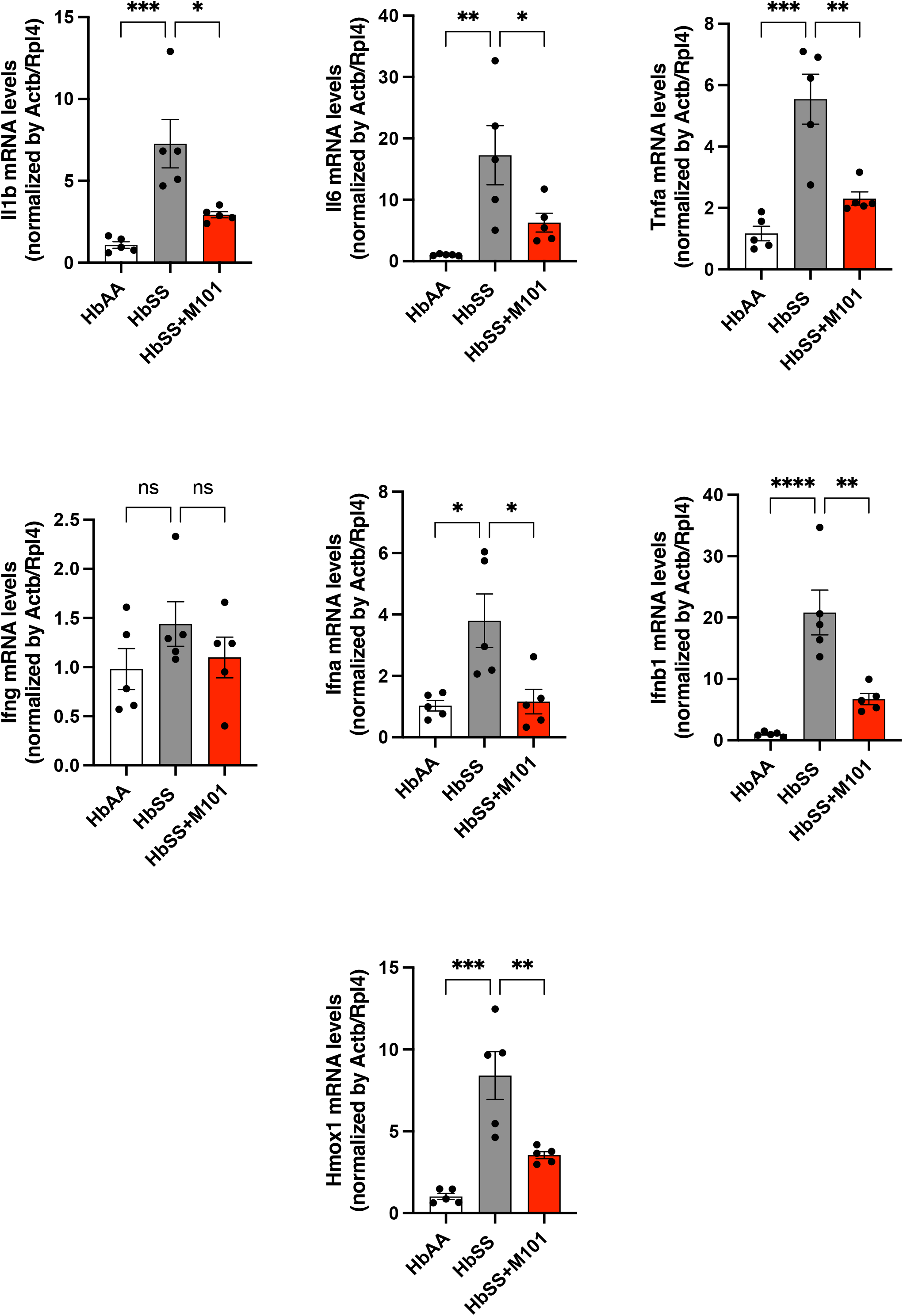
M101 indirectly reduces pulmonary inflammation in hemin-challenged HbSS mice. Expression of Il1b, Il6, Tnfa, Ifna, Ifnb1, Hmox1, and Ifng mRNA in lung tissues from HbSS mice injected with hemin and treated with or without M101 (600 mg/kg). Significant reductions were observed for most inflammatory markers, while Ifng levels remained unchanged. Results are normalized to Actb/Rpl4 and shown as mean ± SEM. p<0.05, p<0.01, p<0.001, p<0.0001.

## Discussion

In this study, we investigated the therapeutic potential of M101, a novel HBOC derived from the marine worm *Arenicola marina*, in a mouse model of SCD. Our findings demonstrate that M101 significantly reduces RBC sickling, mitigates hemolysis, indirectly attenuates inflammation and alleviates microvascular occlusion of lung and liver, under two distinct models of VOC *in vivo*. These results underscore the multifaceted effects of M101 in addressing key pathological features of SCD, particularly tissue hypoxia and oxidative stress, which are major drivers of disease progression.

One of the most notable findings of our study is the potent anti-sickling effect of M101 observed *in vivo*. The dose-dependent reduction in sickled RBCs following M101 administration suggests a direct role in improving oxygen delivery through a physical mechanism and stabilizing RBC morphology under hypoxic conditions. Comparatively, mitapivat, an oral pyruvate kinase (PK) activator, has been shown to improve hemoglobin levels and hemoglobin-oxygen affinity, reducing RBC sickling ^24^. Its mechanism of action involves enhanced RBC glycolysis and ATP production, leading to improved RBC energy metabolism and membrane stability. Clinical trials of mitapivat (e.g., RISE UP trial) demonstrated a significant reduction in hemolysis markers, alongside improved hemoglobin levels ^24,25^. Our results show that M101 reduces hemolysis, as evidenced by decreased LDH

and indirect bilirubin levels, suggesting that M101’s oxygen-carrying and antioxidant properties provide protection against hemolysis and RBC injury. Unlike pyruvate kinase activators, which primarily targets HbS polymerization, M101’s dual function as an oxygen carrier and antioxidant broadens its therapeutic mechanism, allowing it to mitigate both hypoxia and oxidative stress, two interconnected processes central to SCD pathophysiology.

Our study also demonstrates that M101 significantly decreases microvascular occlusion in both hypoxia and heme-induced models of VOC. Mitapivat, while effective in reducing erythrocyte oxidative stress and improving hemoglobin levels *in vivo* ^26^, has yet to demonstrate its ability to reduce vaso-occlusion, particularly during acute events. In contrast, M101’s ability to alleviate microvascular congestion may result from its capacity to improve oxygenation and reduce oxidative stress in hypoxic tissues, thereby preventing endothelial dysfunction and RBC adhesion. This dual effect on oxygen delivery and oxidative stress reduction sets M101 apart as a unique therapeutic candidate for VOC, where hypoxia-driven RBC and endothelial dysfunction exacerbate microvascular obstruction.

The present study reveals that M101 also exerts indirectly, through the correction of hypoxia and oxidative stress, anti-inflammatory and anti-oxidative transcriptional effects in target organs affected during VOC. In both hypoxia and hemin-induced models of SCD, M101 consistently suppressed the expression of pro-inflammatory cytokines such as *Il1b, Il6, Tnfa*, and interferons (*Ifna, Ifnb1, Ifng*), as well as the oxidative stress marker *Hmox1* in the liver and lungs. These findings are particularly relevant given the central role of inflammation and oxidative stress in the amplification of endothelial dysfunction, leukocyte adhesion, and microvascular obstruction in SCD.

Comparatively, existing agents such as hydroxyurea show partial anti-inflammatory benefits. Hydroxyurea reduces leukocyte counts and increases fetal hemoglobin, thereby mitigating downstream inflammation, but does not directly modulate cytokine expression.

In contrast, M101 appears to suppress upstream inflammatory signaling pathways, as indicated by the marked downregulation of key cytokines and type I/II interferons across both liver and lung tissues. The consistent reduction of *Hmox1*, a marker of heme-induced oxidative stress and a sensor of redox imbalance, further supports the antioxidant capacity of M101, which likely derives from its intrinsic SOD-like activity. This dual oxygen-carrying and antioxidative profile distinguishes M101 from classical HBOCs, which have historically failed due to pro-oxidant and vasoactive effects. The ability of M101 to avoid inflammation and oxidative stress positions it as a uniquely comprehensive treatment candidate for SCD—capable of intervening in both the initiation and propagation of vaso-occlusive events, as already demonstrated by RBC transfusion. By mitigating key drivers of endothelial activation and tissue injury, M101 through oxygen transportation holds particular promise not only for preventing acute VOCs but also for protecting against cumulative organ damage, a critical unmet need in the long-term management of SCD.

The antioxidant properties of M101 are particularly noteworthy, as oxidative stress plays a central role in SCD pathology. Hemolysis releases free hemoglobin and heme, which drive the production of ROS, promoting endothelial activation and inflammation. The antioxidant activity may explain the pronounced reduction in vascular congestion observed in M101-treated mice, particularly in the heme-induced VOC model (Figure 8).

**Figure 8.**
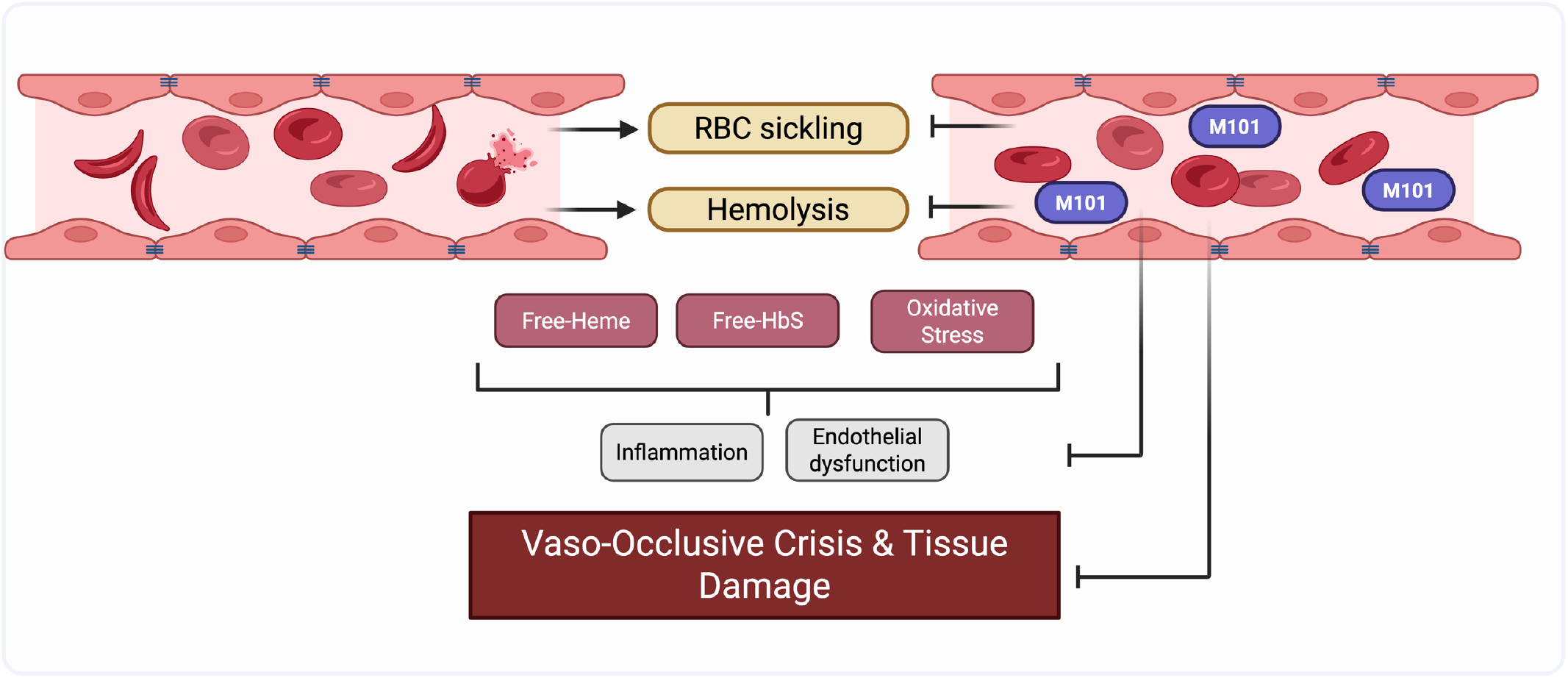
Mechanisms of Vaso-Occlusive Crisis and the Therapeutic Role of M101 in Sickle Cell Disease. Red blood cell (RBC) sickling and hemolysis in sickle cell disease release free heme, free hemoglobin S (HbS), and induce oxidative stress, which collectively trigger inflammation and endothelial dysfunction. These events contribute to the development of vaso-occlusive crises (VOC) and tissue damage. The therapeutic agent M101 mitigates these effects, thereby potentially interrupting the pathological cascade leading to VOC and organ damage.

Red blood cell transfusion has demonstrated clear clinical efficacy in managing or preventing severe acute complications of SCD, such as stroke or acute chest syndrome ^9,27,28^. Chronic transfusion therapy is also the treatment of choice—aside from stem cell transplantation—for secondary stroke prevention, particularly in patients with a history of stroke or elevated transcranial Doppler velocities ^29^. However, the widespread use of transfusions is limited by immunological complications, such as alloimmunization and DHTR; and by infectious risks, especially in regions like Africa. In some cases, DHTR renders further transfusions unsafe ^30^. As a result, oxygen-carrying therapeutics are emerging as a promising alternative, both for acute management and chronic treatment. The preventive effect of M101on organ damage and inflammation observed after a single injection suggests a potential for easy use during acute sickle cell crises. Nevertheless, further studies will be required to assess the long-term efficacy, safety, and pharmacokinetics of M101, particularly in the context of acute VOCs and chronic hemolytic anemia.

In conclusion, our findings highlight the therapeutic potential of M101 as a novel intervention for SCD. By reducing RBC sickling, hemolysis, and microvascular occlusion, M101 demonstrates a unique ability to target key pathological processes in SCD. Future investigations focusing on the clinical translation of M101 will be critical to establishing its role in the therapeutic landscape of SCD.

## Acknowledgment

This work was supported by state funding from the Agence Nationale de la Recherche under the Investissements d’avenir program (ANR-10-IAHU-01).

## Authorship

Contribution: F.Z., L.D., B.B., L.L., O.H., J.B.A and T.T.M. designed the study; R.R.-B. and T.T.M. performed the experiments; F.Z., L.D., B.B., L.L., J.B.A. and T.T.M. analyzed all experiments and drafted the manuscript; and all authors reviewed the manuscript.

## Notes

### Competing Interest Statement

The authors have declared no competing interest.

